# Buffering Agent Induced Lactose Content Increases via Growth Hormone-Mediated Activation of Gluconeogenesis in Lactating Goats

**DOI:** 10.1101/155291

**Authors:** Lin Li, MeiLin He, Ying Liu, Yuanshu Zhang

**Affiliations:** Key Laboratory of Animal Physiology and Biochemistry, Ministry of Agriculture, Nanjing Agricultural University, Nanjing 210095, PR China

**Keywords:** Magnesium oxide, Sodium bicarbonate, high-concentrate diet, Hepatic gluconeogenesis

## Abstract

Dairy goats are often fed a high-concentrate (HC) diet to meet lactation demands; however, long-term concentrate feeding is unhealthy and decreases milk yield and lactose content. Therefore, we tested whether a buffering agent increases the output of glucose in the liver and influences of lactose synthesis. In this study, sixteen lactating goats were randomly assigned to two groups: one group received a HC diets (Concentrate: Forage = 6:4, HG), and the other group received the same diet with a buffering agent added (0.2% NaHCO3, 0.1% MgO, BG) as a treatment for 19-weeks experimental period. The results showed that the total volatile fatty acids and lipopolysaccharide (LPS) declined in the rumen leading to the rumen pH was stabilized in the BG group. Milk yield and lactose content increased. The alanine aminotransferase, aspartate transaminase, alkaline phosphatase, pro-inflammatory cytokines, LPS and lactate content in the plasma was significantly decreased, whereas prolactin and growth hormone levels were increased. The hepatic vein content of glucose was increased. In addition, the expression of pyruvate carboxylase (PC), phosphoenolpyruvate carboxykinase (PEPCK) and glucose-6-phosphatase (G6PC) in the liver was significantly up-regulated. In mammary gland, the glucose transporter type-1, 8, 12 and sodium-glucose cotransporter-1 levels were increased. Cumulatively, the buffering agent treatment increased blood concentrations of glucose via the gluconeogenes and promoting their synthesis in the liver. It may contribute to the increase in milk yield and lactose synthesis of lactating goats.

## Introduction

In the dairy industry, it is currently common practice to feed a high-concentrate (HC) diet to lactating cows or goats to meet their energy requirements to support high milk production. However, long-term high concentrate feeding is harmful to the health of ruminants and leads to a decrease in milk yield [1]. It was reported that feeding of HC diets to lactating cows causes a decline in the rumen pH if organic acids, such as volatile fatty acid (VFA) and lactic acid, accumulate in the rumen[2]. Digestion of a HC diets results in less production of saliva and bicarbonate, and reduced buffering capacity coupled with greater accumulation of organic acids has been reported to increase the incidence of subacute ruminal acidosis (SARA)[3]. A rumen pH of less than 5.8 for over 4 h per day is used as a parameter to determine the occurrence of SARA[4]. In addition, decreased rumen pH results in the release of lipopolysaccharide (LPS), which originate from the cell-wall component of gram-negative bacteria[5]. Previous studies have shown that LPS can translocate into the bloodstream from the digestive tract under conditions of high permeability, and after injury to the liver tissue [6].

In the ruminant, lactose constitutes about 40% of total solids in milk composition. As lactose maintains the osmolarity of milk, the rate of lactose synthesis serves as a major control of the volume of milk yield [7]. Glucose is the main precursor of lactose synthesis in the epithelial cell of the mammary gland, however, the mammary gland cannot synthesize glucose from other precursors due to the lack of glucose 6-phosphatase (G6PC)[8]. Therefore, the mammary gland is dependent on the blood supply for its glucose needs and as a consequence, mammary glucose uptake is a rate-limiting factor for milk yield[9]. Liver glycometabolism of ruminants is different from monogastric animals. In the lactating dairy cows, glucose is supplied primarily by hepatic gluconeogenesis to maintain stable blood glucose[10]. Dairy cows experience an increased demand for glucose to support whole body glucose metabolism and to supply glucose for lactose synthesis[11]. Therefore, liver gluconeogenesis plays an important role in the lactose synthesis of mammary gland.

Buffering agent could enhance the acid base buffer capacity and has been used to prevent ruminant rumen SARA and improve the production performance. Previous studies indicated that the addition of sodium bicarbonate (NaHCO3), magnesium oxide (MgO) to a diet could be given to lactating cows to increase the content of lactose, as well as milk yield[12]. It is well documented that dietary addition of 2% NaHCO3 could increase the buffering capacity and prevent the acidosis in rumen[13].

However, at present, the research of buffering agent is focused on the composition and production of milk from dairy cows. Furthermore, little is known regarding the mechanism of how a buffering agent improves milk yield and lactose content in goats. In this study, we created a buffering agent consisting of (0.2% NaHCO3, 0.1% MgO) and mixed it with a high-concentrate diet source that was fed to lactating goats. We then investigated the effect of this buffering agent on the development of SARA and milk yield and lactose content to elucidate potential mechanisms for this phenomenon.

## Materials and Methods

### Ethical approval

The Institutional Animal Care and Use Committee of Nanjing Agricultural University (Nanjing, People’s Republic of China) approved all of the procedures (surgical procedures and care of goats). The protocol of this study was reviewed and approved specifically, with the project number 2011CB100802. The slaughter and sampling procedures strictly followed the ‘Guidelines on Ethical Treatment of Experimental Animals’(2006) no. 398 set by the Ministry of Science and Technology, China and the ‘Regulation regarding the Management and Treatment of Experimental Animals’ (2008) no. 45 set by the Jiangsu Provincial People’s Government.

### Animals and experimental procedures

Sixteen healthy multiparous mid-lactating saanen goats (body weight, 39 ± 7 kg, mean ± SEM, 3-4 weeks post-partum) at the age of 2-4 years were used in experiments. They were housed in individual stalls in a standard animal feeding house at Nanjing Agricultural University (Nanjing, China). All goats were randomly divided into two groups: one group recieved a high-concentrate diet (Concentrate: Forage = 6:4, HG, n=8), and the other group received the same diet with a buffering agent added (0.2% NaHCO3, 0.1% MgO, purchased from Nanjing Jiancheng Bioengineering Institute, China, BG, n=8) as a treatment. The ingredients and nutritional composition of the diets are presented in Table 1. The animals were fed the respective diets for 19 weeks, and had free access to water during the experimental period.

**Table 1.** Composition and nutrient levels of experimental diets

| Concentrate : Forage ratio 6 : 4 |  |
| --- | --- |
| Ingredient (%) |  |
| Leymus chinensis | 27.00 |
| Alfalfa silage | 13.00 |
| Corn | 23.24 |
| Wheat bran | 20.77 |
| Soybean meal | 13.67 |
| Limestone | 1.42 |
| NacL | 0.30 |
| Premix<sup>a</sup> | 0.60 |
| Total | 100.00 |
| Nutrient levels<sup>b</sup> |  |
| Net energy/(MJ.kg-1) | 6.71 |
| Crude protein/% | 16.92 |
| Neutral detergent fiber/% | 31.45 |
| Acid detergent fiber/% | 17.56 |
| Calcium/% | 0.89 |
| Phosphorus/% | 0.46 |

a. Provided per kg of diet: VA 6000IU/kg, VD 2500IU/kg, VE 80mg/kg, Cu 6.25 mg/kg, Fe 62.5 mg/kg, Zn 62.5 mg/kg, Mn 50mg/kg, I 0.125 mg/kg, Co 0.125 mg/kg.
b. Nutrient levels were according to National Research Council (NRC,2001).

Prior to the initiation of the experiment, all of the goats were installed rumen fistula and hepatic catheters. Specific steps are as follows: First, all goats were installed rumen cannula, and indwelt with hepatic and portal vein catheters, under general anaesthesia by inhalation of isoflurane (2.5% in 1:1 mixture of oxygen and air; Abbott Scandinavia AB, Solna, Sweden). Reflexes (pupillary and palpebral) and respiratory rate were monitored before and during the surgery to verify anaesthetic depth. Next, a small cannula constructed from polyvinylchloride was inserted into the rumen of each goat[14]. After rumen fistula surgery, all goats were ruminally cannulated, fitted with indwelling catheters in the portal vein and hepatic vein[15]. At the end of these procedures the goats received an intramuscular injection of flunixin meglumine [2 mg (kg body weight)^−1^], an anti-inflammatory and analgesic (Banamine; Mantecorp Ind. Quím. e Farm. Ltda, Rio de Janeiro, Brazil) and their respiratory movements were monitored until they regained consciousness. Goats were continuously monitored for 1 h after the surgery, and all animals used in the experiments on the day after presented in a good and stable clinical condition with no signs of inflammation or pain problems. After surgery, goats were observed for 2 weeks during recovery from the surgery. Sterilized heparin saline (500 IU/mL, 0.3 mL/time) was administered at 6-hour intervals every day until the end of the experiment to prevent catheters from becoming blocked.

### Rumen fluid collection and analysis

Fifteen minutes prior to feed delivery and 0, 2, 4, 6, 8 and 10 h after feed delivery on 7 consecutive days during week 19, 20 mL rumen fluids was collected with a nylon bag and the pH value was measured immediately with pH-meter.

The rumen fluid was collected and each sample was transferred into a 50-mL sterile tube and kept on ice until transported to the laboratory for the initial processing before LPS determination. Another part of each rumen fluid sample was centrifuged at 3,200 × g for 10 min at 4°C immediately after collection and the supernatant was collected. To analyze VFA in ruminal fluid, a 5-mL aliquot was deproteinized with 1 mL of 25% metaphosphoric acid. These samples were stored at -20°C until analysis.

The concentration of LPS in rumen fluid was measured by a Chromogenic End-point Tachypleus Amebocyte Lysate Assay Kit (Chinese Horseshoe Crab Reagent Manufactory Co. Ltd, Xiamen, China). Pretreated rumen fluid samples were diluted until their LPS concentrations were in the range of 0.1-1.0 endotoxin units (EU)/mL relative to the reference endotoxin.

VFA were measured using capillary column gas chromatography (GC-14B, Shimadzu, Japan; Capillary Column: 30 m × 0.32 mm × 0.25 mm film thickness; Column temperature = 110°C, injector temperature = 180°C, detector temperature = 180°C).

### Plasma biochemical parameters analysis

In the 19^th^ week, blood samples were collected from the jugular vein, hepatic vein and portal vein blood in 10 mL vacuum tubes containing sodium heparin. Blood was centrifuged at 3000 x *g* for 15 min to separate the plasma, which was then stored at -20°C until analysis. The plasma glucose content was quantified using a Beckman Kurt AU5800 series automatic biochemical analyzer (Beckman Kurt, USA) at the General Hospital of Nanjing Military Region (Nanjing, China).

The growth hormone (GH), tumor necrosis factor-a (TNF-a), interleukin 1β (IL-1β) concentration in the plasma were measured by radioimmunoassay with commercially available human radioimmunoassay kits purchased from Beijing North Institute of Biological Technology. The detection range of radioimmunoassay kits for GH (rabbit, B12PZA), TNF-a (rabbit, C06PZA) and IL-1β (rabbit, C09PDA) were 0.1-50 ng/mL, 1-10 ng/mL and 0.1-8.1 ng/mL, respectively. All of the procedures were performed according to the manufacturer’s instructions.

The analyses for the prolactin, glucocorticoids, histamine and lactate were performed using Enzyme-Linked Immunosorbent Assay (ELISA) kit (Shanghai Enzyme-linked Biotechnology Co. Ltd, Shanghai, China) according to the manufacturer’s instructions. The detected range of ELISA kits for prolactin, glucocorticoids, histamine and lactate were 5-2000 pg/mL, 0-80 ng/mL, 2-600 ng/mL and 0.1-30 mmol/mL, respectively. The LPS concentration were determined using a chromogenic endpoint assay (CE64406, Chinese Horseshoe Crab Reagent Manufactory Co., Ltd., Xiamen, China) with a minimum detection limit of 0.01 EU/mL. The procedures were performed according to the manufacturer’s instructions.

### Milk composition analysis

Goats were milked at 8:30 h and 18:30 h, and the milk yield was recorded daily. A 50-mL milk sample was taken to determine the lactose content once a week (Milk-Testing™ Milkoscan 4000, FOSS, Denmark) at the Animal Experiment Center of College of Animal Science and Technology at the Nanjing Agricultural University.

### Sample collection

In the 19^th^ week, mammary gland tissues were obtained by biopsy after 4 h after the morning feeding. Local anesthesia (2% lidocaine hydrochloride) was administered into breast skin in a circular pattern surrounding the incision site, then a 2cm incision was made and mammary gland tissue was dissected. Tissue samples (500-800 mg) were rinsed with 0.9% saline, snap frozen in liquid nitrogen and were used for RNA extraction. Goats were slaughtered after overnight fasting. Incisions were sutured, and antibiotics were administered intramuscularly to avoid infection.

After 19 weeks, all goats were killed with neck vein injections of xylazine [0.5 mg (kg body weight)^−1^; Xylosol; Ogris Pharme, Wels, Austria] and pentobarbital [50 mg (kg body weight)^−1^; Release; WDT, Garbsen, Germany]. After slaughter, liver tissue was collected and washed twice with cold physiological saline (0.9% NaCl) to remove blood. Livers were then transferred into liquid nitrogen and used for RNA and protein extraction.

### RNA isolation, cDNA synthesis and real-time PCR

Total RNA was extracted from liver samples using TRIzol reagent (15596026, Invitrogen, USA) and converted to cDNA using commercial kits (Vazyme, Nanjing, China). All PCR primers were synthesized by Generay Company (Shanghai, China); the primer sequences are listed in Table 2. PCR was performed using the AceQ qPCR SYBR Green Master Mix Kit (Vazyme, Nanjing, China) and the MyiQ2 Real-time PCR System (Bio-Rad, USA) with the following cycling conditions: 95°C for 2 min, 40 cycles of 95°C for 15 sec and 60°C for 30 sec. Glyeraldehyde 3-phosphate dehydrogenase (GAPDH) served as a reference for normalization. The _2_^−⎕⎕Ct^ method was used to analyse the real-time PCR results, and each gene mRNA level is expressed as the fold change relative to the mean value of the control group.

**Table 2.**
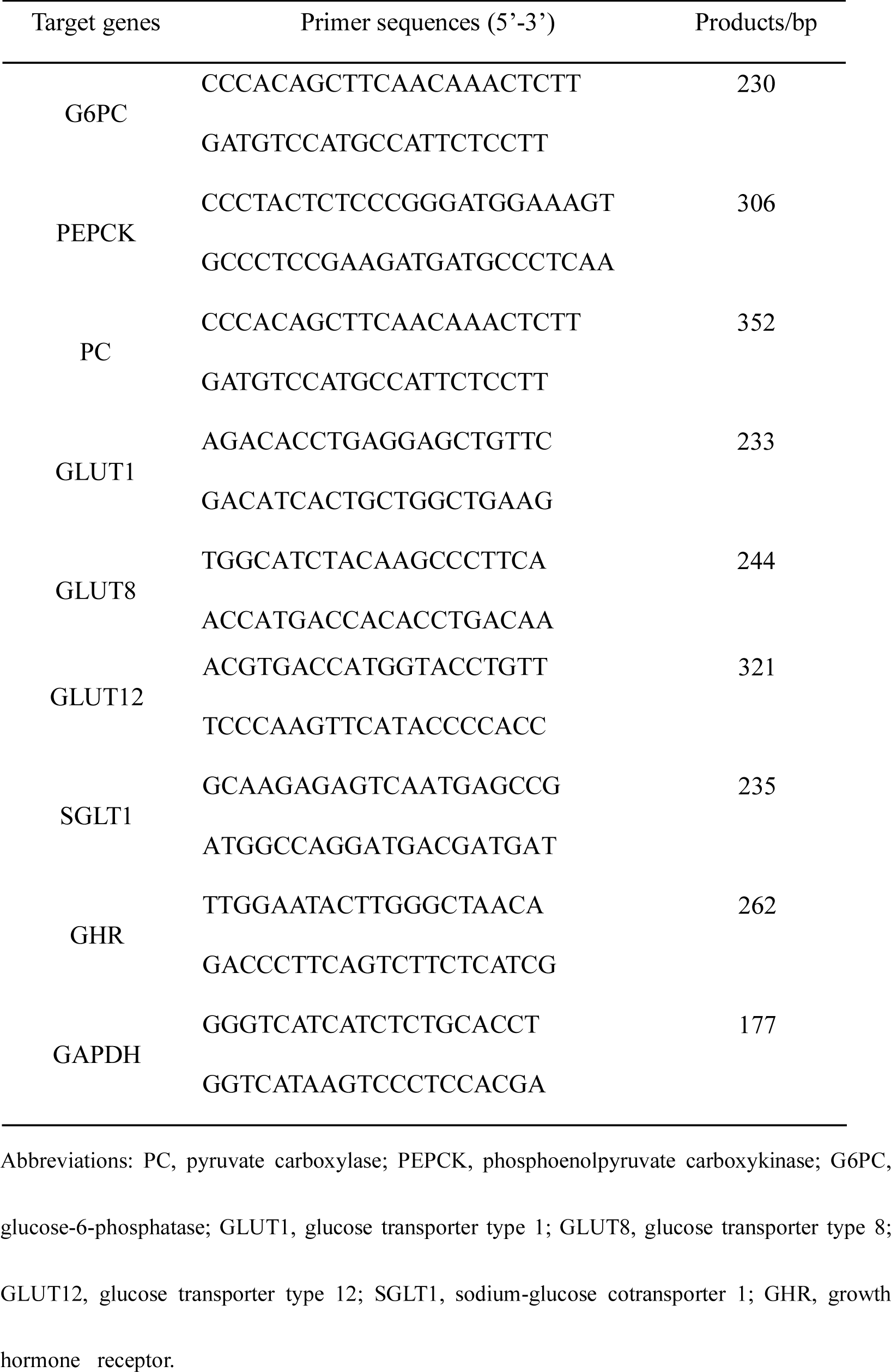
Primer sequences and product size

### Western blot analysis

Total protein was extracted from frozen liver samples, and the concentration was determined using a bicinchoninic acid (BCA) assay kit (Pierce, Rockford, IL, USA). We isolated 30 μg of total protein from each sample, which was subjected to electrophoresis on a 10% SEMS-PAGE gel. The separated proteins were transferred onto nitrocellulose membranes (Bio Trace, Pall Co., USA). The blots were incubated with the following primary antibodies for overnight at 4°C at dilutions of 1:1000 in block: rb-anti-phosphoenolpyruvate carboxykinase (rb-anti-PEPCK, #12940, CST), rb-anti-glucose transporter type 1 (rb-anti-GLUT1, ab14683, Abacm) and rb-anti-glucose transporter type 12 (rb-anti-GLUT12, ab100993, Abacm). A rb-anti-GAPDH primary antibody (A531, Bioworld, China, 1:10,000) was also incubated with the blots to provide a reference for normalization. After washing the membranes, an incubation with HRP-conjugated secondary antibody was performed for 2 h at room temperature. Finally, the blots were washed, and the signal was detected by enhanced chemiluminescence (ECL) using the LumiGlo substrate (Super Signal West Pico Trial Kit, Pierce, USA). The ECL signal was recorded using an imaging system (Bio-Rad, USA) and analyzed with Quantity One software (Bio-Rad, USA).

### Statistical analysis

The results were expressed as mean ± SEM. The data of ruminal pH and glucose in plasma of hepatic vein, portal vein and jugular vein were analyzed for differences due to diet, feeding time, and their interactions by Univariate using the General Linear Models of SPSS 11.0 for Windows (StatSoft Inc, Tulsa, OK, USA). The differences in milk yield, lactose content, plasma biochemical index, mRNA and protein expression between two groups were analyzed by Independent-Samples T test using the Compare Means of SPASS 11.0 for Windows (StaSoft Inc, Tulsa, OK, USA). Data were considered statistically significant if *P* < 0.05, *P* < 0.01. The numbers of replicates used for statistics are noted in the Tables and Figures.

## Results

### The buffering agent treatment increased daily milk yield and lactose content in lactating goats

We quantified the daily milk yield and lactose content in the milk of the two experimental groups from 1 to 19 weeks of treatment. During week 1 to week 2, there was no significant difference in average daily milk yield and lactose content between BG goats and HG goats. However, the average daily milk yield (*P* < 0.05) and lactose content (*P* < 0.01) increased significantly in the BG group from 3-19 weeks of treatment compared to the HG group (Fig. 1).

**Figure 1.**
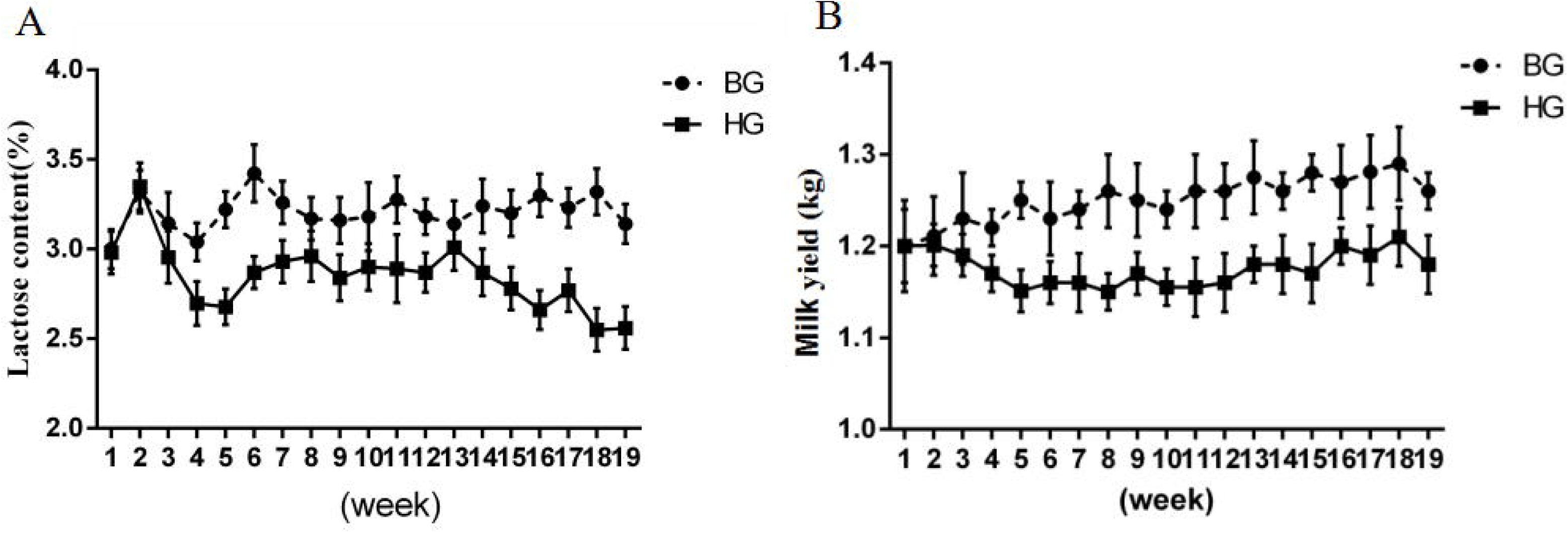
Comparison of the average weekly lactose content and milk yield between the buffering agent (BG) and high-concentrate diet (HG) groups. Values are shown as means ± SEM, n = 8/group. \**P* < 0.05 compared with the HG group.

### The buffering agent treatment stabilized ruminal fluid pH in lactating goats fed a high-concentrate diet

After feeding 19 weeks, the dynamic pH curve in the BG group was higher than that in the HG group during the long-term experiment. It showed that a pH value under 5.8 lasted for 6 h in the HG group, which indicated that SARA was successfully induced. The pH value of the BG group was significantly increased in comparison to those in the HG group (*P* < 0.05). However, ruminal pH was affected significantly by digesting time, while there was no interaction of digesting time and diet on ruminal pH (*P* > 0.05; Fig. 2).

**Figure 2.**
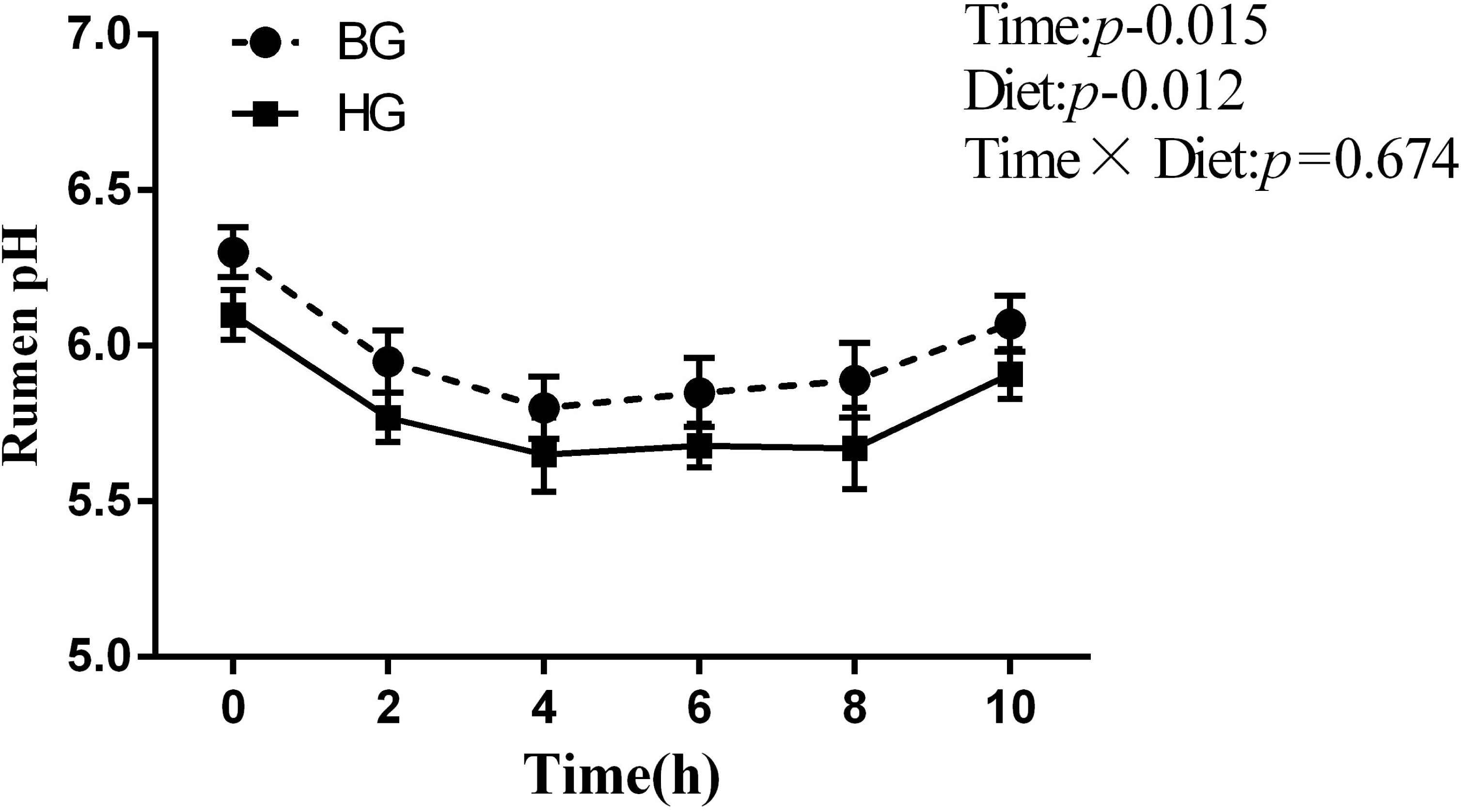
pH value in ruminal fluid after 19 weeks feeding regime. Data were analyzed for differences due to diet, time, and their interactions by Univariate using the General Linear Models of SPSS 11.0 for Windows (StatSoft Inc, Tulsa, OK, USA). Values are mean ± SEM, n = 8/group. \**P* < 0.05 compared with the HG group.

### VFA and LPS concentrations in rumen fluid

As shown in Table 3, BG goats showed significantly lower level of LPS concentration in rumen fluid than HG goats. Concentrations of total VFA, propionate, butyrate in rumen fluid were significantly decreased in BG goats compared to HG (*P* < 0.05). However, the ratio of propionate to butyrate in the rumen was significantly elevated in the BG group (*P* < 0.05).

**Table 3.**
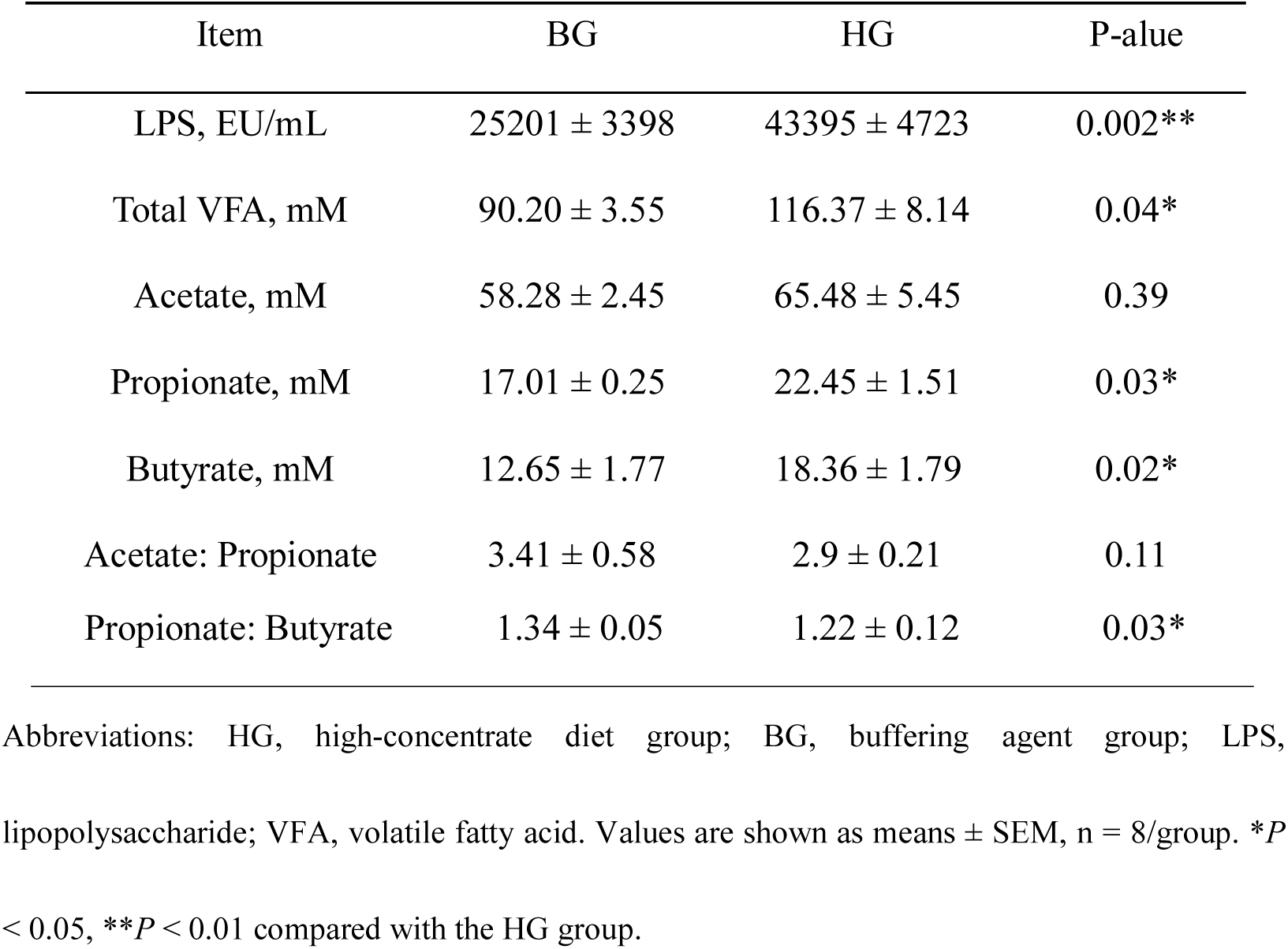
Effects of the buffering agent treatment on rumen fermentation parameters in goats

### The buffering agent treatment changed plasma hormone, enzyme, primary pro-inflammatory cytokines and metabolic produced in the lactating goats

As shown in Table 4, the plasma content of alanine aminotransferase (ALT), aspartate transaminase (AST) and alkaline phosphatase (AKP) were significantly lower in the BG group compared to the HG group (*P* < 0.05). Although the plasma content of lactic dehydrogenase (LDH) was declined, there was no significant difference between BG and HG groups. The pro-inflammatory cytokines including TNF-a and IL-1β in the BG were significantly lower than that in the HG goats (*P* < 0.05). Meanwhile, we found that the metabolism products of LPS, histamine and lactate content were also lower in the BG goats compared to the HG goats. Among them, LPS and lactate were significant different (*P* < 0.05). Moreover, the BG goats showed significantly higher levels of GH and prolactin concentration in plasma than HG goats, while there was no significant difference of glucocorticoids concentration in plasma between BG and HG goats.

**Table 4.** Effects of the buffering agent treatment on plasma enzyme, primary pro-inflammatory cytokines, metabolic produced and hormone of lactating goats

| Item | BG | HG | P-value |
| --- | --- | --- | --- |
| Plasma biochemical parameter |  |  |  |
| ALT (IU/L) | $40.33 \pm 2.84$ | $77.67 \pm 10.44$ | 0.03* |
| AST (IU/L) | $43.33 \pm 4.48$ | $71.33 \pm 8.67$ | 0.04* |
| LDH (IU/L) | $233.66 \pm 16.45$ | $243.66 \pm 13.54$ | 0.66 |
| AKP (IU/L) | $109.67 \pm 17.07$ | $213.5 \pm 20.50$ | 0.02* |
| TNF-α (ng/mL) | $2.47 \pm 0.30$ | $4.61 \pm 0.48^*$ | 0.03* |
| IL-1β (ng/mL) | $0.74 \pm 0.03$ | $0.79 \pm 0.05$ | 0.04* |
| LPS (EU/mL) | $1.82 \pm 0.14$ | $3.76 \pm 1.13$ | 0.04* |
| Histamine (ng/mL) | $1.99 \pm 0.06$ | $2.11 \pm 0.09$ | 0.09 |
| Lactate (mmol/L) | $0.95 \pm 0.05$ | $1.39 \pm 0.16^*$ | 0.04* |
| Hormone levels |  |  |  |
| Prolactin (pg/mL) | $436.57 \pm 37.78$ | $373.29 \pm 30.59$ | 0.04* |
| Glucocorticoids (ng/mL) | $10.2 \pm 1.67$ | $9.8 \pm 2.56$ | 0.08 |
| Growth hormone (ng/mL) | $0.94 \pm 0.08$ | $0.67 \pm 0.03$ | 0.03* |

Abbreviations: BG, buffering agent group; HG, high-concentrate diet group; ALT, alanine aminotransferase; AST, aspartate transaminase; LDH, lactic dehydrogenase; AKP, alkaline
phosphatase; TNF-a, tumor necrosis factor-a; IL-1β , interleukin 1β ; LPS, lipopolysaccharide;
Values are shown as means ± SEM, n = 8/group. \* $P$ < 0.05 compared with the HG group.

### The buffering agent treatment regulated enzymes required for glucose transfer in the mammary gland of lactating goats

We found that the mRNA expression of glucose transporter type 1 (GLUT1), glucose transporter type 8 (GLUT 8), glucose transporter type 12 (GLUT12) and sodium-glucose cotransporter 1 (SGLT1) were also higher in the BG goats compared to expression in the HG goats. In particular, expression of GLUT1 and SGLT1 were significantly higher than that in the HG goats (*P* < 0.05). The level of GLUT1 protein expression in the mammary gland was significantly up-regulated in BG goats compared to HG (*P* < 0.05). Additionally, there was a tendency increase in protein expression of GLUT12 in BG goats (Fig. 3).

**Figure 3.**
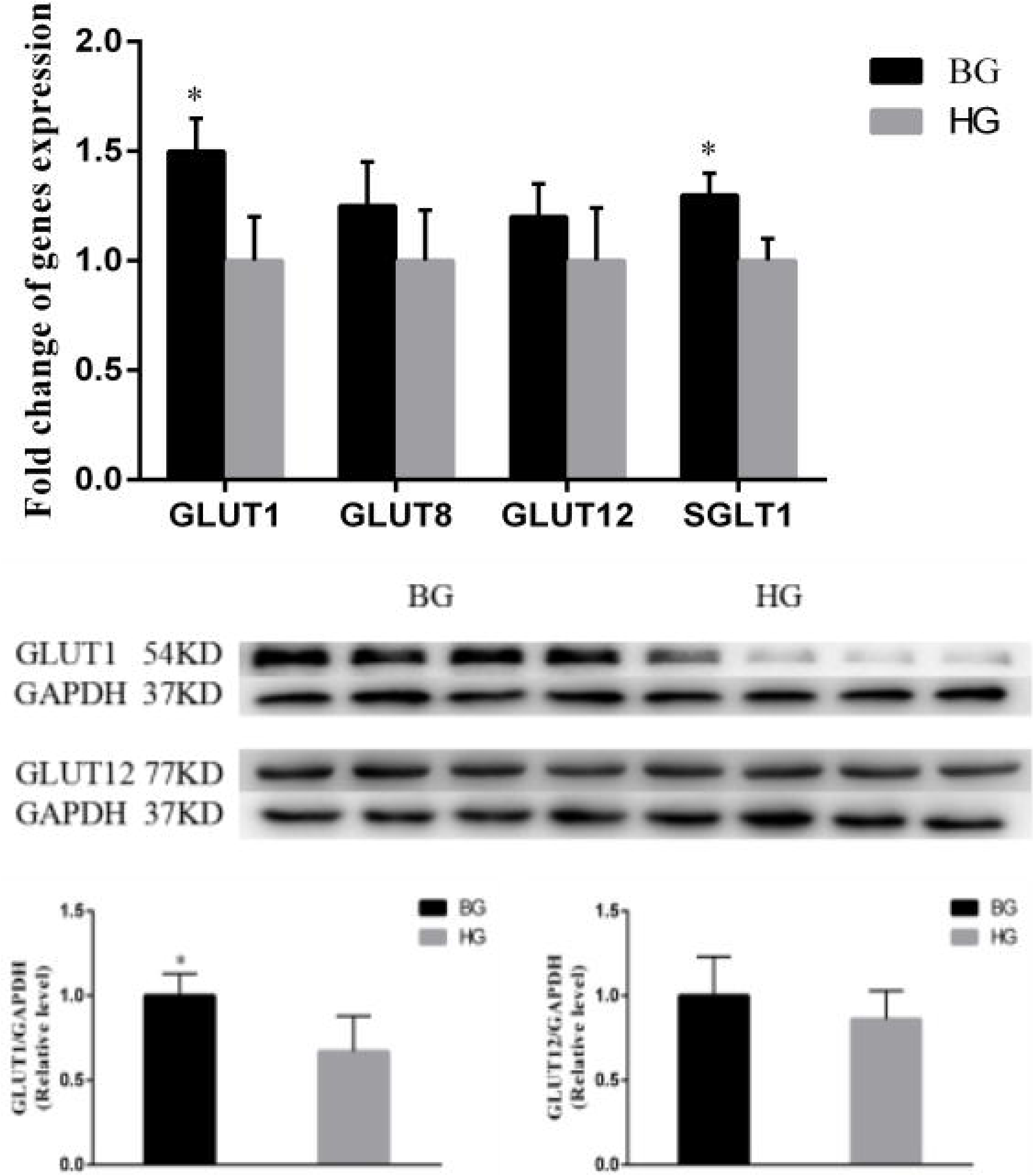
Effects of buffering agent treatment on the expression of mammary gland glucose transfer genes in lactating goats. The experiments were repeated three times. Values are shown as means ± SEM, n = 3. \**P* < 0.05 compared with the HG group.

### The buffering agent treatment increased production of glucose in the liver

We next examined glucose in plasma obtained from the jugular vein, hepatic vein and portal vein of both treatment groups. After 19 weeks feeding, the jugular and hepatic vein content of glucose was significantly increased in the BG group compared to the HG group (*P* < 0.01). The portal vein content of glucose was increased, but there was no significant difference between BG and HG groups. Compared to HG group, we found that the glucose content of BG group was significantly higher in the hepatic vein compared to the portal vein (*P* < 0.05, Table 5). This indicates that more glucose is produced in the liver. It is possible that the synthesis of glucose was activated following treatment with the buffering agent.

**Table 5.** The average concentrations of glucose in plasma of hepatic vein, portal vein and jugular vein of lactating goats

| Glucose (mmol/L) | BG | HG | Effect, p-value |  |  |
| --- | --- | --- | --- | --- | --- |
| | | | Diet | Time | Diet × Time |
| Hepatic vein |  |  |  |  |  |
| 0h | 3.34 ± 0.37* | 3.01 ± 0.18 | 0.003 | 0.292 | 0.636 |
| 4h | 3.35 ± 0.37* | 3.15 ± 0.18 | | | |
| 8h | 3.44 ± 0.37* | 3.07 ± 0.18 | | | |
| Portal vein |  |  |  |  |  |
| 0h | 3.27 ± 0.11 | 3.26 ± 0.13 | 0.102 | 0.902 | 0.494 |
| 4h | 3.28 ± 0.12 | 3.27 ± 0.12 | | | |
| 8h | 3.27 ± 0.09 | 3.25 ± 0.15 | | | |
| Jugular vein |  |  |  |  |  |
| 0h | 3.30 ± 0.05 | 3.27 ± 0.09 | 0.002 | 0.890 | 0.579 |
| 4h | 3.33 ± 0.24 | 3.29 ± 0.12 | | | |
| 8h | 3.34 ± 0.14 | 3.25 ± 0.04 | | | |

Abbreviations: HG, high-concentrate diet group; BG, buffering agent group. Values are shown as
means ± SEM, n = 8/group. \* $P$ < 0.05 compared with the HG group.

### The buffering agent treatment regulated enzymes required for gluconeogenesis and GHR in the livers of lactating goats

We found that the mRNA expression of phosphoenolpyruvate carboxykinase (PEPCK) and pyruvate carboxylase (PC), glucose-6-phosphatase (G6PC) was higher in the BG goats compared to expression in the HG goats. In particular, expression of PEPCK and G6PC were significantly higher than that in the HG goats (*P* < 0.05). The level of PEPCK protein expression in the liver was significantly up-regulated in BG goats compared to HG (*P* < 0.05). This is consistent with our previous observation that PEPCK mRNA expression increases in BG goats (Fig. 4). Taken together, these results suggested that treatment with the buffering agent promoted the gluconeogenesis in the liver. The level of GHR expression in the liver was significantly up-regulated in BG goats compared to HG (*P* < 0.05, Fig. 5).

**Figure 4.**
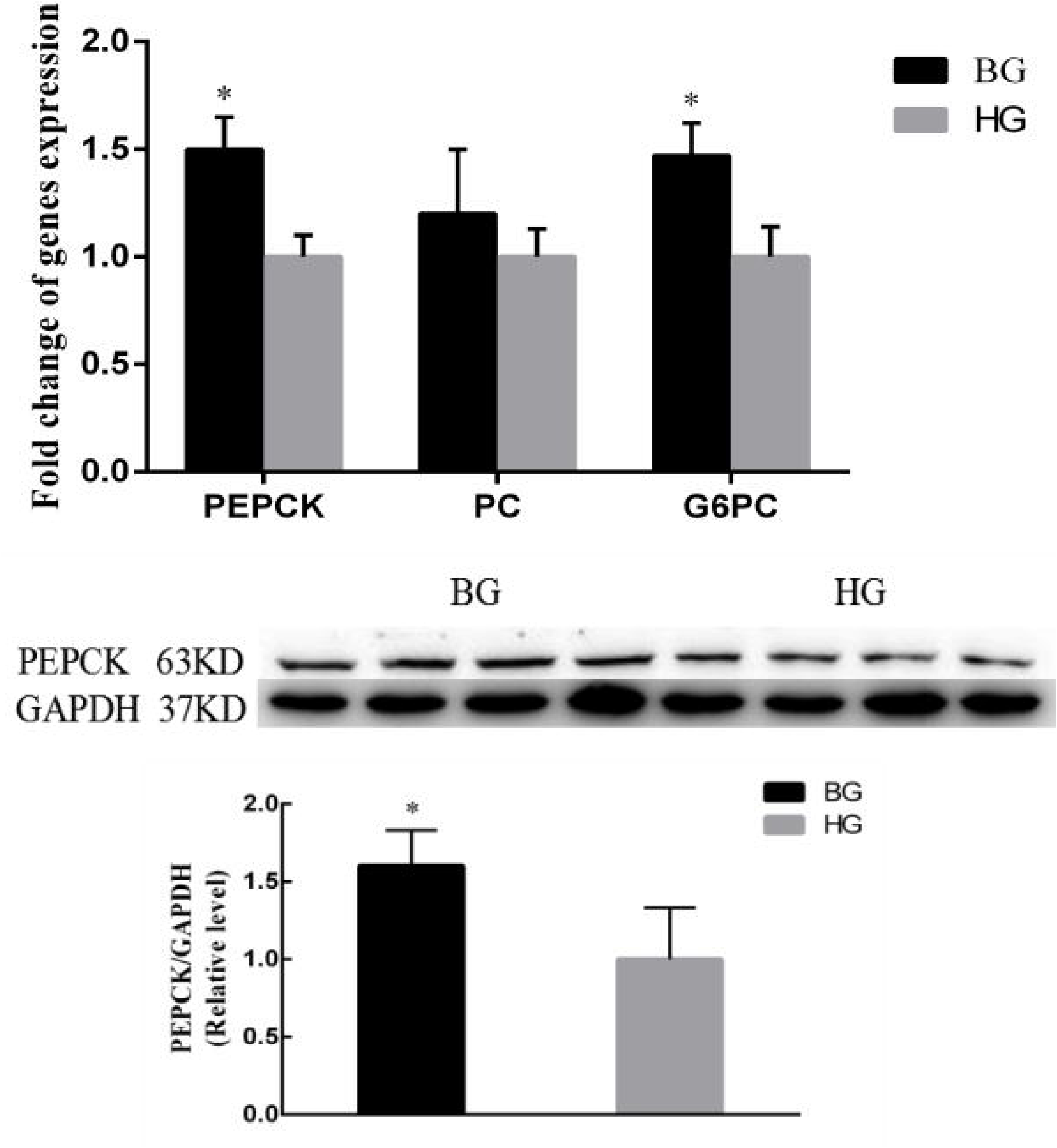
Effects of buffering agent treatment on the expression of liver gluconeogenesis genes in lactating goats. The experiments were repeated three times. Values are shown as means ± SEM, n = 3. \**P* < 0.05, compared with the HG group.

**Figure 5.**
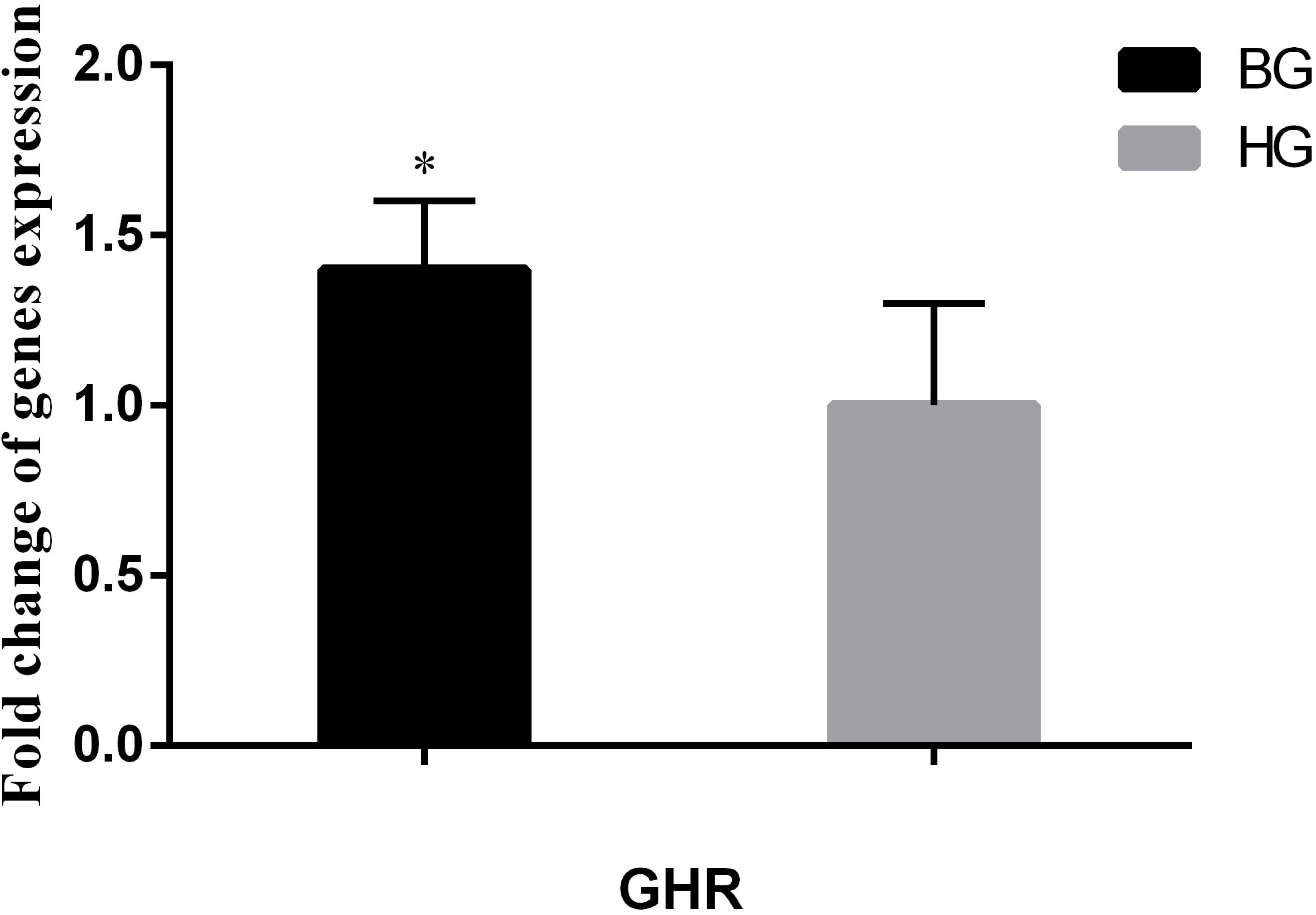
Effects of buffering agent treatment on the expression of GHR in livers of lactating goats. The experiments were repeated three times. Values are shown as means ± SEM, n = 3. \**P* < 0.05 compared with the HG group.

**Figure 6.**
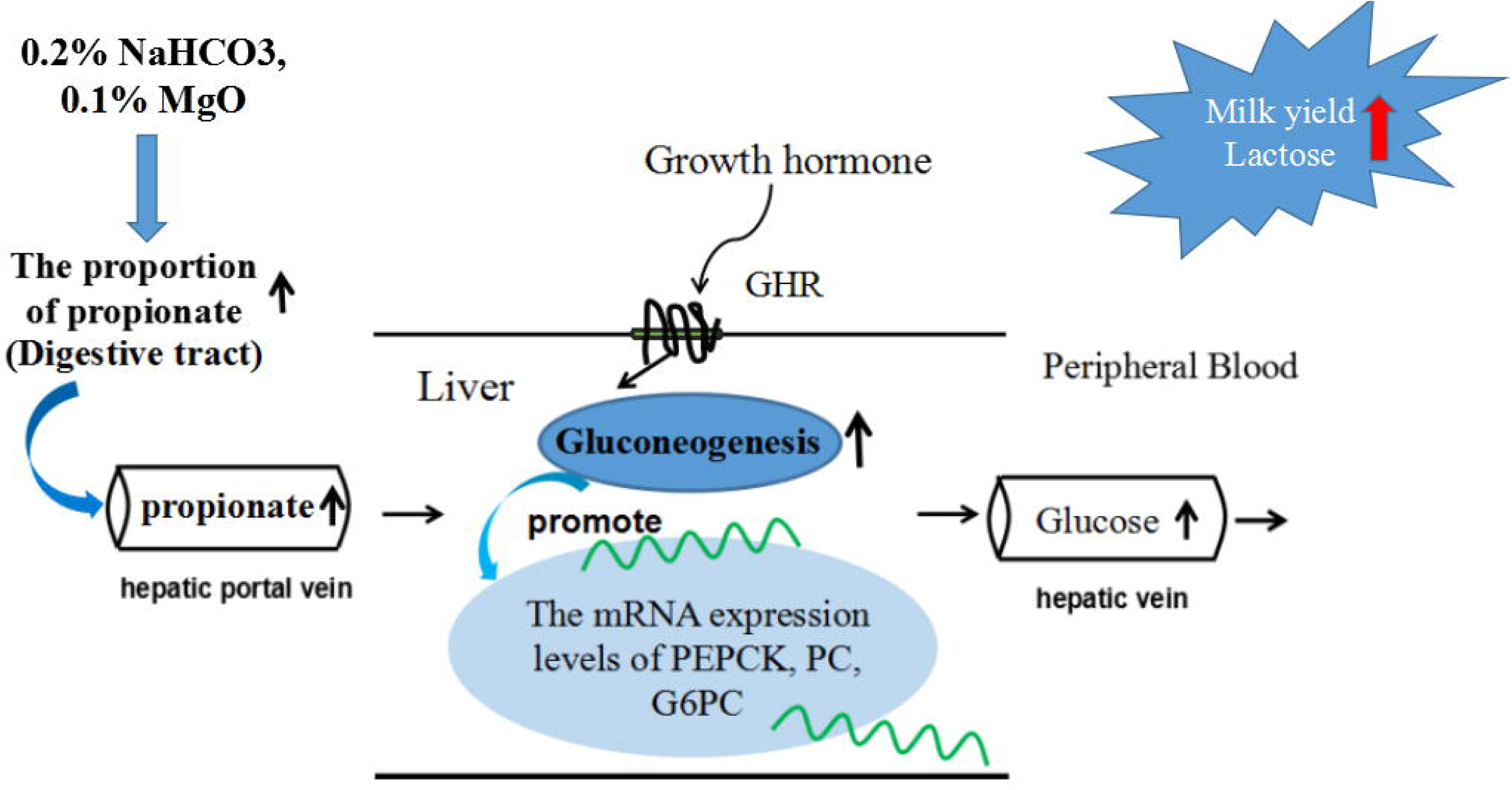
Growth hormone activates the gluconeogenesis to regulate glucose synthesis in the liver. Abbreviations: PEPCK, phosphoenolpyruvate Carboxykinase; PC, pyruvate carboxylase; G6PC, glucose-6-phosphatase.

## Discussion

In recent years, dairy goats are often fed HC diets to meet the energy demand for high milk yields. However, the consumption of a HC diets is harmful to the health of dairy goats [16,17]. It’s well documented that feeding HC diets to ruminants results in subacute ruminal acidosis (SARA), a common metabolic disease especially occurred in high-producing animals. The root cause is that excessive amounts of rapidly fermentable nonstructural carbohydrates increase the accumulation of organic acids and shift of microbial population in gastrointestinal tract in ruminants [18]. Moreover, an increased amount of fermentable carbohydrates, such as starch, pass through the forestomach to the intestinal tract in acidosis, which accelerates intestinal tract fermentation[19]. It ultimately affects the intestinal absorption of nutrients. Importantly, in accordance with previous research, long-term feeding a high-concentrate diet could induce the depression of the content of lactose and milk yield[20].

The NaHCO3 could increase the buffering capacity and prevent acidosis in the rumen. It was reported that the rumen pH profile improved and a higher yield of milk and milk solids when NaHCO3 was supplemented to a high-concentrate diet [21]. Previous studies indicated that the addition of NaHCO3 and MgO to restricted-roughage rations for goats could increase the content of lactose and milk yield [22]. Prolactin is involved in the development of the mammary gland, and the start and continuation of lactation by influencing lactogenesis. It is responsible for the synthesis of the lactose and milk production found in milk [23]. In our experiment, the duration of a rumen pH less than 5.8 lasted for 4h in the goats fed a high-concentrate diet. According to the definition of experimental SARA, HG goats were suffering from SARA disease. However, after feeding 19 weeks, the buffering agent added to the high-concentrate diet stabilized ruminal pH and prevented the occurrence of SARA. Meanwhile, a increase in the milk yield and lactose content was observed in the BG group. The concentrations of prolactin in blood was also markedly increased. Therefore, its increased levels in the blood are associated with milk yield and lactose content improvement.

It is well known that the feeding of HC diets leads to the translocation of LPS from gram-negative bacteria in the gastrointestinal tract into the circulating blood. Other studies showed that feeding a diet containing 60% concentrate to lactating goats elevated blood LPS concentrations[24]. The increased levels of circulating LPS also can elevated the concentration of blood pro-inflammatory cytokines IL-1β and TNF-α and activation of liver inflammatory responses [25,26]. The biochemical parameters ALT, AST and AKP in peripheral blood are common indicators used to assess the status of liver function^27^. In particular, ALT is a specific parameter that reflects hepatocyte damage. In the present study, we observed that feeding of a HC diets induced the massive release of LPS in the rumen, which could trigger a local or systemic inflammatory response after the translocation of LPS into the bloodstream. Furthermore, our data demonstrated that the feeding of a HC diets significantly increased the concentrations of LPS, TNF-α and IL-1β in the plasma. The increase in blood pro-inflammatory cytokines is consistent with the translocation of LPS and activation of inflammatory responses. In addition, the concentrations of ALT, ALP and AKP in peripheral blood were also higher in the HG goats compared to the BG goats. These results provide insight into feeding the HC diets resulted in the breach of hepatocytes releasing these enzymes into the circulation. Importantly, the results showed that those pro-inflammatory cytokines, LPS, TNF-α and IL-1β in the plasma of the BG goats were significantly lower than that in the HG goats. Therefore, we hypothesized that the buffering agent added to the high-concentrate diet reduced the release of the rumen LPS and stabilized the body health of lactating goats.

Compared to monogastric animals, glucose is supplied primarily by hepatic gluconeogenesis to maintain stable blood glucose content in the ruminants^10^. Therefore, the liver plays a crucial physiological role in the body, and is responsible for glucose metabolism. Our study documented that feeding HC diet to lactating goats for a long time leads to the LPS-cytokines-induced the inflammatory response, and increases the consumption and catabolism of glucose in liver [28]. GH is a polypeptide hormone synthesized and secreted by the anterior pituitary gland, which plays a key role in regulating ruminant mammary gland development and lactation [29]. It is important for regulating glycometabolism, in part through its promotion of gluconeogenesis in the liver [30]. The body’s health is essential to the normal production of hormones. However, the increased translocation of LPS may into the brain via the blood enhances the inflammatory response, which might be ultimately affect on the levels of the growth hormone. PEPCK and G6PC are two key hepatic gluconeogenic enzymes the expression and activity of which are increased hepatic glucose output [31]. PC is the first regulatory enzyme in gluconeogenic pathway that converts pyruvate to oxaloacetate in gluconeogenesis [32]. Major glucose precursors of ruminant liver include propionate, amino acids and lactate. It has been documented that the increased proportion of propionate may be related to glycogenesis in ruminant. Because most VFA emerge in the portal vein after absorption from the digestive tract [33], alterations in proportion of propionate influence the gluconeogenesis in the liver. Therefore, the liver gluconeogenesis plays a crucial physiological role in maintaining the body blood sugar levels, as it is the main organ for glucose storage, in the form of glycogen, as well as endogenous glucose production [34]. Our results indicated that the buffering agent added to the high-concentrate diet significantly decreased the total VFA, propionate and butyrate levels in the ruminal fluid. However, the ratio of propionate to butyrate was increased in the BG group. We also observed that the buffering agent treatment promoted the expression of PEPCK, PC and G6PC, indicating that gluconeogenesis in the liver was increased. In addition, the BG diets increased glucose content in hepatic vein. The plasma GH and GHR levels were also increased in the BG goats: elevated GH increases the glucose content and activity of gluconeogenesis in the liver. Meanwhile, the buffering agent added to the HC diet inhibited the consumption of glucose and stabilized the liver health of lactating goats. Taken together, these findings suggest that the feeding of BG diets can promote liver gluconeogenesis by the increased proportion of propionate in the rumen, and the increased entry of glucose into the blood through the hepatic vein.

In lactating animals, providing glucose for the mammary gland is a metabolic priority because glucose is the primary precursor for lactose synthesis in the mammary gland. Once taken up by the lactating mammary epithelial cell, glucose is either used in the synthesis of lactose or processed by glycolysis to provide energy. Lactose is synthesized from free glucose and uridine diphosphate (UDP)-galactose by lactose synthase catalysis [35]. The mammary gland itself cannot synthesize glucose from other precursors because of the lack of glucose-6-phosphatase [8,36]. Therefore, the mammary gland is dependent on the blood supply for its glucose needs. In addition, lactose maintains the osmolarity of milk, the rate of lactose synthesis serves as a major factor influencing milk yield. It is also indicated that lactose synthesis and milk yield show a linear or positive correlation with glucose uptake in the mammary gland of goats and cows [37,38]. Glucose uptake in the mammary gland is increased dramatically during lactation. As is known, glucose transport across the plasma membranes of mammalian cells is carried out by 2 distinct processes: facilitative transport, mediated by a family of facilitative glucose transporters (GLUT); and sodium-dependent transport, mediated by the Na+/glucose cotransporters (SGLT) [39]. An early study demonstrated that facilitated GLUT 1, GLUT 8, GLUT 12 and SGLT1 have different expression in mammary gland [40]. The GLUT1 is ubiquitously expressed in lactating cow tissues, being most abundant in the mammary gland and kidney and lowest in the omental fat and skeletal muscle [41]. SGLT1 play a important role in glucose transport of Golgi membrane [42]. In our experiment, we found that the glucose content in the plasma of the jugular vein was increased in the BG group compared to the HG group. GLUT 1, GLUT 8, GLUT 12 and SGLT1 expression in mammary gland were also elevated in the BG goats. Moreover, the level of GLUT 1 protein was significantly enhanced in mammary gland of BG goats. Taken together, it is indicated that the buffering agent added to the high-concentrate diet led to the translocation of more glucose from the peripheral blood into the mammary epithelial cells and thereby increased the milk yield and lactose content.

## Conclusions

In summary, we systematically investigated the effects of the buffering agent on milk quality of lactating goats and found milk yield and lactose content were increased. Furthermore, the blood GH and prolactin levels were increased in the BG goats: elevated GH increases the hepatic gluconeogenesis and activity. Activated gluconeogenesis promotes levels of blood glucose released from the liver (Figure 9). Thus, the increased glucose in the hepatic vein during of goats fed a BG diets may be play a key role in increasing the milk yield and lactose synthesis of lactating goats. However, the GLUT1, 8, 12 and SGLT1 in mammary gland were also elevated in BG goats. It is possible that the buffering agent added to the HC diet inhibited the release of inflammatory cytokine and stabilized the mammary gland health of lactating goats. Then it causes to the increase in glucose transporters in the mammary gland and prolactin levels in the blood, which could also increase the lactose content in milk. Therefore, further research is needed to elucidate the underlying mechanism.

## Author contributions

L.L. performed the experiment and drafted the manuscript. M.H., and Y.L. performed the experiment and analysed the data. Y.Z. contributed to experimental design and manuscript revision. L.L., and Y.Z. conceived the idea, designed the experiment and finalized the manuscript. All authors read and approved the final manuscript.

## Acknowledgements

All authors thank Xiangzhen Shen (Department of clinical veterinary, Nanjing Agricultural University) for technical help during animal experimentation.

## Compliance with Ethical Standards

The study was approved by the ethical committee of Nanjing agricultural university.

## References

(1) Xu T, Tao H, Chang G, Zhang K, Xu L & Shen X (2015). Lipopolysaccharide derived from the rumen down-regulates stearoyl-CoA desaturase 1 expression and alters fatty acid composition in the liver of dairy cows fed a high-concentrate diet. BMC veterinary research 11,

(2) Chen Y & Oba M (2012). Variation of bacterial communities and expression of Toll-like receptor genes in the rumen of steers differing in susceptibility to subacute ruminal acidosis. Vet Microbiol 159, 451–459.

(3) Emmanuel DG, Dunn SM, Ametaj BN. Feeding high proportions of barley grain stimulates an inflammatory response in dairy cows. J Dairy Sci. 2008;91(2):606–14.

(4) Beauchemin KA, Yang WZ & Rode LM (2003). Effects of particle size of alfalfa-based dairy cow diets on chewing activity, ruminal fermentation, and milk production. J Dairy Sci 86, 630–643.

(5) Tao S, Tian J, Cong R, Sun L, Duanmu Y, Dong H, Ni Y & Zhao R (2015). Activation of cellular apoptosis in the caecal epithelium is associated with increased oxidative reactions in lactating goats after feeding a high-concentrate diet. Exp Physiol 100, 278–287.

(6) Duanmu Y, Cong R, Tao S, Tian J, Dong H, Zhang Y, Ni Y & Zhao R (2016). Comparative proteomic analysis of the effects of high-concentrate diet on the hepatic metabolism and inflammatory response in lactating dairy goats. J Anim Sci Biotechnol 7, 1.

(7) Neville MC, Allen JC & Watters C (1983). The mechanisms of milk secretion. Lactation. Springer US 49, 102.

(8) Threadgold LC & Kuhn NJ (1979). Glucose-6-phosphate hydrolysis by lactating rat mammary gland. Int J Biochem 10, 683–685.

(9) Kronfeld DS (1982). Major metabolic determinants of milk volume, mammary efficiency, and spontaneous ketosis in dairy cows. J Dairy Sci 65, 2204–2212.

(10) Reynolds CK (2006). Production and metabolic effects of site of starch digestion in dairy cattle. Anim. Feed Sci. Technol 130, 78–94.

(11) Li WQ, Bu DP, Wang JQ, Nan XM, Sun P & Zhou LY (2013). Effect of two different diets on liver gene expression associated with glucose metabolism in dairy cows. Livest Sci 158, 223–229.

(12) Lingxin D & Runhou Z (2000). Effects of supplementing by-pass protein and buffer additives in the diet of lactating cows on milk output and composition. Xibei Nonglin Keji Daxue Xuebao (China).

(13) Islam SMS, Hossain MA, Hashim MMA, Sarker MSA & Paul AK (2014). Effects of sodium bicarbonate on induced lactic acidosis in Black Bengal goats. Wayamba J Anim Sci 6, 1044–1057.

(14) Stoddard GE, Allen NN, Hale WH, Pope AL, Sorensen DK & Winchester WR (1951) A permanent rumen fistula cannula for cows and sheep. J Anim Sci 10, 417–423.

(15) Huntington GB, Reynolds CK & Stroud BH (1989). Techniques for measuring blood flow in splanchnic tissues of cattle. J Dairy Sci 72, 1583–1595.

(16) Tao S, Han Z, Tian J, Cong R, Duanmu Y, Dong H, Ni Y & Zhao R (2016). Downregulation of prostaglandin E2 is involved in hindgut mucosal damage in lactating goats fed a high-concentrate diet. Exp Physiol 101, 272–281.

(17) Li L, Cao Y, Xie Z, Zhang Y (2017). A High-Concentrate Diet Induced Milk Fat Decline via Glucagon-Mediated Activation of AMP-Activated Protein Kinase in Dairy Cows. Sci Rep 7, 44217.

(18) Plaizier JC, Krause DO, Gozho GN & McBride BW (2008). Subacute ruminal acidosis in dairy cows: The physiological causes, incidence and consequences. The Vet J 176, 21–31.

(19) Li S, Khafipour E, Krause DO, Kroeker A, Rodriguez-Lecompte JC, Gozho GN & Plaizier JC (2012). Effects of subacute ruminal acidosis challenges on fermentation and endotoxins in the rumen and hindgut of dairy cows. J Dairy Sci 95, 294 303.

(20) Chang G, Zhang K, Xu T, Jin D, Seyfert HM, Shen X & Zhuang S (2015). Feeding a high-grain diet reduces the percentage of LPS clearance and enhances immune gene expression in goat liver. BMC Vet Res 11, 1.

(21) Cruywagen CW, Taylor S, Beya MM & Calitz T (2015). The effect of buffering dairy cow diets with limestone, calcareous marine algae, or sodium bicarbonate on ruminal pH profiles, production responses, and rumen fermentation. J Dairy Sci 98, 5506–5514.

(22) Lee MJ & Hsu AL (1991). Effect of supplementation of sodium bicarbonate and magnesium oxide in diet on lactating performance and rumen characteristics of dairy goats. J Chin Soc Anim Sci 20, 431–442.

(23) Alipanah M, Kalashnikova L & Rodionov G (2007). Association of prolactin gene variants with milk production traits in Russian Red Pied cattle. Iran J Biotechnol 5, 158–161.

(24) Dong H, Wang S, Jia Y, Ni Y, Zhang Y, Zhuang S, Shen X & Zhao R (2013). Long-term effects of subacute ruminal acidosis (SARA) on milk quality and hepatic gene expression in lactating goats fed a high-concentrate diet. PLoS One 8, e82850.

(25) Chang G, Zhuang S, Seyfert HM, Zhang K, Xu T, Jin D, Guo J & Shen X (2015). Hepatic TLR4 signaling is activated by LPS from digestive tract during SARA, and epigenetic mechanisms contribute to enforced TLR4 expression. Oncotarget 6, 38578.

(26) Duanmu Y, Cong R, Tao S, Tian J, Dong H, Zhang Y, Ni Y & Zhao R (2016). Comparative proteomic analysis of the effects of high-concentrate diet on the hepatic metabolism and inflammatory response in lactating dairy goats. J Anim Sci Biotechnol 7, 5.

(27) Sevinc M, Basoglu A, Birdane FM & Boydak M (2001). Liver function in dairy cows with fatty liver. Revue de Medecine Veterinaire (France).

(28) Jiang X, Zeng T, Zhang S & Zhang Y (2013). Comparative proteomic and bioinformatic analysis of the effects of a high-grain diet on the hepatic metabolism in lactating dairy goats. PloS one 8, e80698.

(29) Akers RM (2006). Major advances associated with hormone and growth factor regulation of mammary growth and lactation in dairy cows. J Dairy Sci 89, 1222–1234.

(30) Emmison N, Agius L & Zammit VA (1991). Regulation of fatty acid metabolism and gluconeogenesis by growth hormone and insulin in sheep hepatocyte cultures. Effects of lactation and pregnancy. Biochem J 274, 21–26.

(31) Lochhead PA, Salt IP, Walker KS, Hardie DG & Sutherland C (2000). 5-aminoimidazole-4-carboxamide riboside mimics the effects of insulin on the expression of the 2 key gluconeogenic genes PEPCK and glucose-6-phosphatase. Diabetes 49, 896–903.

(32) Pershing RA, Moore SEM, Dinges AC, Thatcher WW & Badinga L (2002). Short communication: hepatic gene expression for gluconeogenic enzymes in lactating dairy cows treated with bovine somatotropin. J Dairy Sci 85, 504–506.

(33) Bergman EN (1990). Energy contributions of volatile fatty acids from the gastrointestinal tract in various species. Physiol Rev 70, 567–590.

(34) Sharabi K, Tavares CD, Rines AK & Puigserver P (2015). Molecular pathophysiology of hepatic glucose production. Mol Aspects Med 46, 21–33.

(35) Watkins WM & Hassid WZ (1962). Synthesis of Lactose by Particulate Enzyme Preparations from Guinea Pig and Bovine Mammary Glands. Sci 136, 329.

(36) Scott RA, Bauman DE & Clark JH (1976). Cellular gluconeogenesis by lactating bovine mammary tissue. J Dairy Sci 59, 50–56.

(37) Cant JP, Trout DR, Qiao F & Purdie NG (2002). Milk synthetic response of the bovine mammary gland to an increase in the local concentration of arterial glucose. J Dairy Sci 85, 494–503.

(38) Huhtanen P, Vanhatalo A & Varvikko T (2002). Effects of abomasal infusions of histidine, glucose, and leucine on milk production and plasma metabolites of dairy cows fed grass silage diets. J Dairy Sci 85, 204–216.

(39) Zhao FQ & Keating AF (2007). Functional properties and genomics of glucose transporters. Curr Genomics 8, 113–128.

(40) Zhao FQ & Keating AF (2007). Expression and regulation of glucose transporters in the bovine mammary gland. J Dairy Sci 90, E76–E86.

(41) Zhao FQ, Glimm DR & Kennelly JJ (1993). Distribution of mammalian facilitative glucose transporter messenger RNA in bovine tissues. Int J Biochem 25, 1897–1903.

(42) Faulkner A, Chaiyabutr N, Peaker M, Carrick DT & Kuhn NJ (1981). Metabolic significance of milk glucose. J Dairy Res 48, 51–56.

